# 8-Oxoguanine transcriptional miscoding can transiently reproduce PAH-associated premature-stop transcripts

**DOI:** 10.64898/2026.10.01.756055

**Authors:** Santiago Moreno-Caceres, Alexis Hayes, Amber Delmas Eliason, David P. Marciano

**Affiliations:** Herbert Wertheim College of Medicine, Florida International University

**Keywords:** 8-oxoguanine, transcriptional mutagenesis, pulmonary arterial hypertension, *BMPR2*, haploinsufficiency, nonsense-mediated decay

## Abstract

Pulmonary arterial hypertension (PAH) is associated with increased oxidative DNA damage, while haploinsufficiency of *BMPR2* and other dosage-sensitive genes contributes directly to disease. Template-strand 8-oxoguanine (8-oxoG) provides a potential transient route to the same loss-of-function outcome through transcriptional mutagenesis without altering genomic DNA. RNA polymerase II can insert adenine opposite the lesion and generate a coding-sense C>A error in the nascent transcript. We asked whether this chemistry could generate premature-stop transcripts in dosage-sensitive PAH genes. Across 17,880 C>A-accessible coding positions in pulmonary vascular genes, 1,192 could generate premature termination codons, including 883 predicted to engage nonsense-mediated decay (NMD). Restricting these sites to expressed autosomal-dominant PAH genes with definitive ClinGen validity and established haploinsufficiency identified 116 NMD-prone positions in *BMPR2, KDR, SMAD9*, and *TBX4*. Eight exactly reproduce ClinVar pathogenic or likely pathogenic stop variants, including five in *BMPR2*. At *BMPR2* c.1424, template-strand 8-oxoG and the expert-panel-reviewed genomic c.1424C>A allele is predicted to generate the same UAA-containing transcript. Published oxidized-guanine maps detect signal at seven of the eight coordinates, providing coordinate-level support for oxidative modification at these sites. These findings define a sequence-resolved route through which repairable oxidative DNA damage could transiently reproduce loss-of-function transcripts encoded by pathogenic PAH alleles without permanent genomic mutation.

## 1 Introduction

Oxidative DNA damage has been documented in pulmonary hypertension and in PAH vascular cells. Oxidized guanine has been detected in lungs from patients with severe pulmonary hypertension [1], and pulmonary artery endothelial cells from patients with PAH show increased reactive-oxygen-species-associated DNA damage that antioxidant treatment reduces [2]. Hypoxic signaling can also generate sequence-localized oxidative guanine lesions in pulmonary endothelial DNA [3]. These observations place oxidative DNA damage within the pulmonary vascular disease environment but do not establish whether individual lesions transiently alter coding output before repair.

When 8-oxoguanine (8-oxoG) occurs on the template strand, RNA polymerase II can insert adenine opposite the lesion and generate a coding-sense C>A change in the nascent RNA while genomic DNA remains unchanged [4–6]. Structural and biochemical studies explain how this mispair escapes RNA polymerase II fidelity checkpoints [7,8], and site-specific experiments have shown that the chemistry generates functional mutant transcripts whose abundance increases when lesion repair is impaired [9,10]. The altered sequence exists only in transcripts synthesized while the damaged template persists, which separates transcriptional miscoding from a stable genomic mutation.

This chemistry is particularly relevant to PAH because haploinsufficiency of *BMPR2* and other dosage-sensitive genes contributes directly to disease [11,12]. A premature termination codon can reduce functional output further by directing the affected RNA into nonsense-mediated decay [13]. A transient transcription error that introduces such a codon could temporarily reproduce the consequence of a pathogenic truncating allele.

We therefore asked whether the sequence constraints of 8-oxoG transcriptional miscoding include positions at which transient C>A errors could generate premature-stop transcripts in dosage-sensitive PAH genes. We mapped these sites across pulmonary vascular genes and then applied PAH gene-disease validity, inheritance, haploinsufficiency, and vascular-expression criteria to identify disease-relevant candidates. We next asked where the transient RNA sequence exactly matches that produced by a pathogenic genomic stop variant, which identifies *BMPR2* c.1424 as the clearest case.

## 2 Results

### 2.1 8-Oxoguanine miscoding can create predicted NMD-prone premature stops in PAH haploinsufficiency genes

Across the pulmonary vascular gene panel, template-strand 8-oxoG could generate 1,192 premature termination codons, of which 883 are predicted NMD-prone under the primary transcript-architecture rule. Restricting these sites to expressed autosomal-dominant genes with definitive PAH gene-disease validity and established haploinsufficiency identified 116 positions in *BMPR2, KDR, SMAD9* and *TBX4* (Figure 1). Each is a coding-strand cytosine paired with a template-strand guanine; oxidation would introduce a premature stop in a dosage-sensitive PAH gene, reducing functional output and contributing to disease.

**Figure 1.**
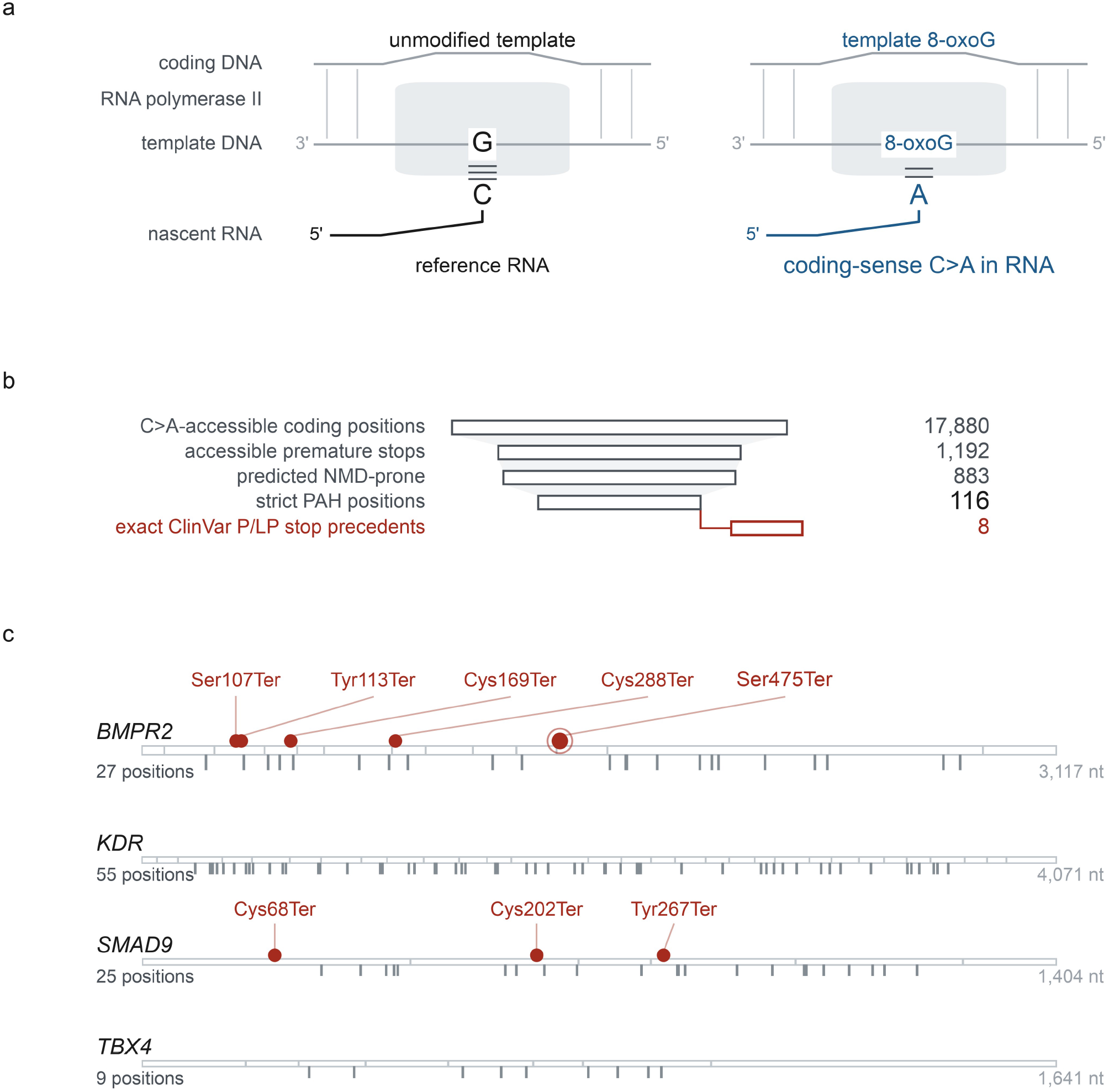
8-Oxoguanine transcriptional miscoding can access premature-stop sequences in PAH haploinsufficiency genes. Throughout, blue marks transient 8-oxoG miscoding and red marks the permanent genomic allele. (**a**) Template-strand 8-oxoG directs adenine incorporation by RNA polymerase II, producing a coding-sense C>A change in RNA while genomic DNA is unaltered. (**b**) Successive biological filters narrow the panel to the strict PAH candidate set, with exact pathogenic or likely pathogenic genomic precedents drawn as a separate branch rather than a further filter. Funnel widths are scaled to log10(count). (**c**) Coding-coordinate maps of *BMPR2, KDR, SMAD9* and *TBX4*. Gray ticks are candidate positions; red lollipops are exact genomic precedents, and the outer ring marks *BMPR2* c.1424. Introns are omitted. P/LP denotes pathogenic or likely pathogenic, NMD denotes nonsense-mediated decay, and Ter denotes translation termination.

*BMPR2* contributes 27 positions, *KDR* 55, *SMAD9* 25, and *TBX4* 9, distributed across their coding regions rather than concentrated within a single local sequence context (Figure 1c). The primary surveillance call is made in the MANE Select transcript, and the set is not uniformly NMD-prone across isoforms: 51 positions are predicted NMD-prone in every GENCODE-primary transcript containing the site, 8 are predicted to escape in at least one such transcript, and 57 occur in a single primary transcript (Supplementary Figure S1). The strict set remained stable across the tested surveillance thresholds, ranging from 113 to 121 positions; all eight exact genomic precedents were retained under the 150-nucleotide and 200-nucleotide start-proximal rules, while seven remained at 250 nucleotides (Supplementary Table S2). The complete 116-position set is unchanged under both the extended and the core vascular-expression definitions at the 5 percent and 10 percent detection thresholds (Supplementary Table S3).

### 2.2 Eight 8-oxoG-accessible stop codons exactly match ClinVar pathogenic or likely pathogenic PAH stop variants

Eight of the 116 candidate positions reproduce a ClinVar pathogenic or likely pathogenic premature-stop variant exactly. Five occur in *BMPR2* and three in *SMAD9*, where the transient C>A transcription error and the permanent genomic C>A variant generate the same premature-stop RNA sequence (Figure 2). Six are predicted NMD-prone in every primary transcript containing the site under the primary classification rule. These records lack equivalent clinical evidence because their ClinVar review levels range from no assertion criteria to expert-panel review, and *BMPR2* c.1424C>A, p.Ser475Ter, is the strongest genetic precedent because it is the only one reviewed by an expert panel. These records establish the pathogenicity of genomic alleles that encode these stop-containing transcripts, while 8-oxoG miscoding provides a transient route to the same RNA sequence without altering genomic DNA.

**Figure 2.**
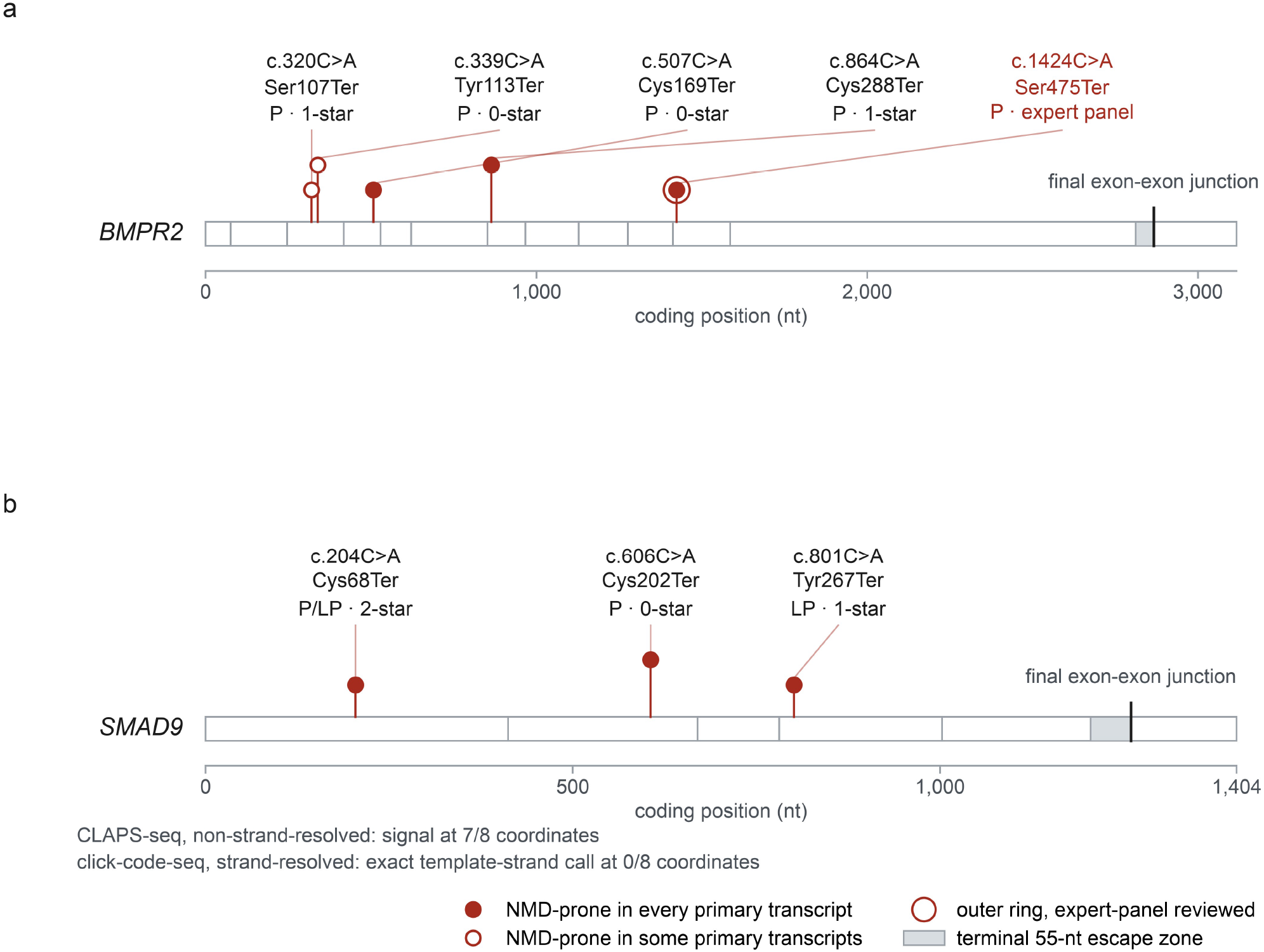
Eight 8-oxoG-accessible stop codons exactly match ClinVar pathogenic or likely pathogenic PAH stop variants. Coding positions of five *BMPR2* (**a**) and three *SMAD9* (**b**) genomic C>A stop variants on the MANE Select coding architecture, with stem heights alternating only to separate closely spaced positions. Filled red markers are predicted NMD-prone in every GENCODE-primary transcript containing the site and open red markers are isoform-dependent. The outer ring identifies *BMPR2* p.Ser475Ter and the shaded interval is the terminal 55-nucleotide NMD escape zone. Each record carries its ClinVar review level, which the marker color does not encode: 0-star for no assertion criteria, 1-star for criteria from a single submitter, 2-star for criteria from multiple submitters without conflict, and expert panel for review by a ClinGen variant curation expert panel. The two lines below the panel’s report lesion-map support separately for each assay, because CLAPS-seq is not strand resolved while click-code-seq is. P denotes pathogenic, LP likely pathogenic, P/LP a record classified as either, NMD nonsense-mediated decay and Ter translation termination.

The archived ClinVar release preceding the analysis snapshot contained 136 pathogenic or likely pathogenic premature-stop single-nucleotide variants across the four strict genes. Nine are coding C>A variants, eight of which fall within the predicted NMD-prone set, while the ninth lies in a terminal exon and is predicted to escape surveillance. This comparison is descriptive and does not test whether C>A-reachable positions are over-represented among reported variants.

Non-strand-resolved CLAPS-seq shows endogenous oxidized-guanine signal at seven of the eight coordinates, including *BMPR2* c.1424 in two of three replicates [14]. Because these tracks report normalized coverage rather than strand-resolved lesion calls, they do not establish the template-strand configuration required for miscoding. None of the eight sites carries an exact template-strand call in the available strand-resolved click-code-seq dataset [15], and both resources derive from nonvascular cells. The maps support oxidative modification at these coordinates while leaving template-strand lesion formation in pulmonary vascular cells unresolved. Per-site values are in Supplementary Table S5.

### 2.3 *BMPR2* c.1424 defines an experimentally tractable convergence site

*BMPR2* c.1424 provides a direct sequence-level convergence between the transient lesion-driven route and a pathogenic genomic allele. The genomic c.1424C>A allele encodes p.Ser475Ter and is classified as pathogenic by the ClinGen pulmonary hypertension variant curation expert panel. Oxidation of the corresponding template guanine to 8-oxoG, followed by adenine incorporation during transcription, would produce the identical UAA stop codon while the genomic sequence remains wild type. In the MANE Select transcript, the resulting stop lies 1,441 coding nucleotides upstream of the final exon-exon junction and is predicted NMD-prone, with the same classification retained in the second GENCODE-primary transcript containing the site (Figure 3a,b).

**Figure 3.**
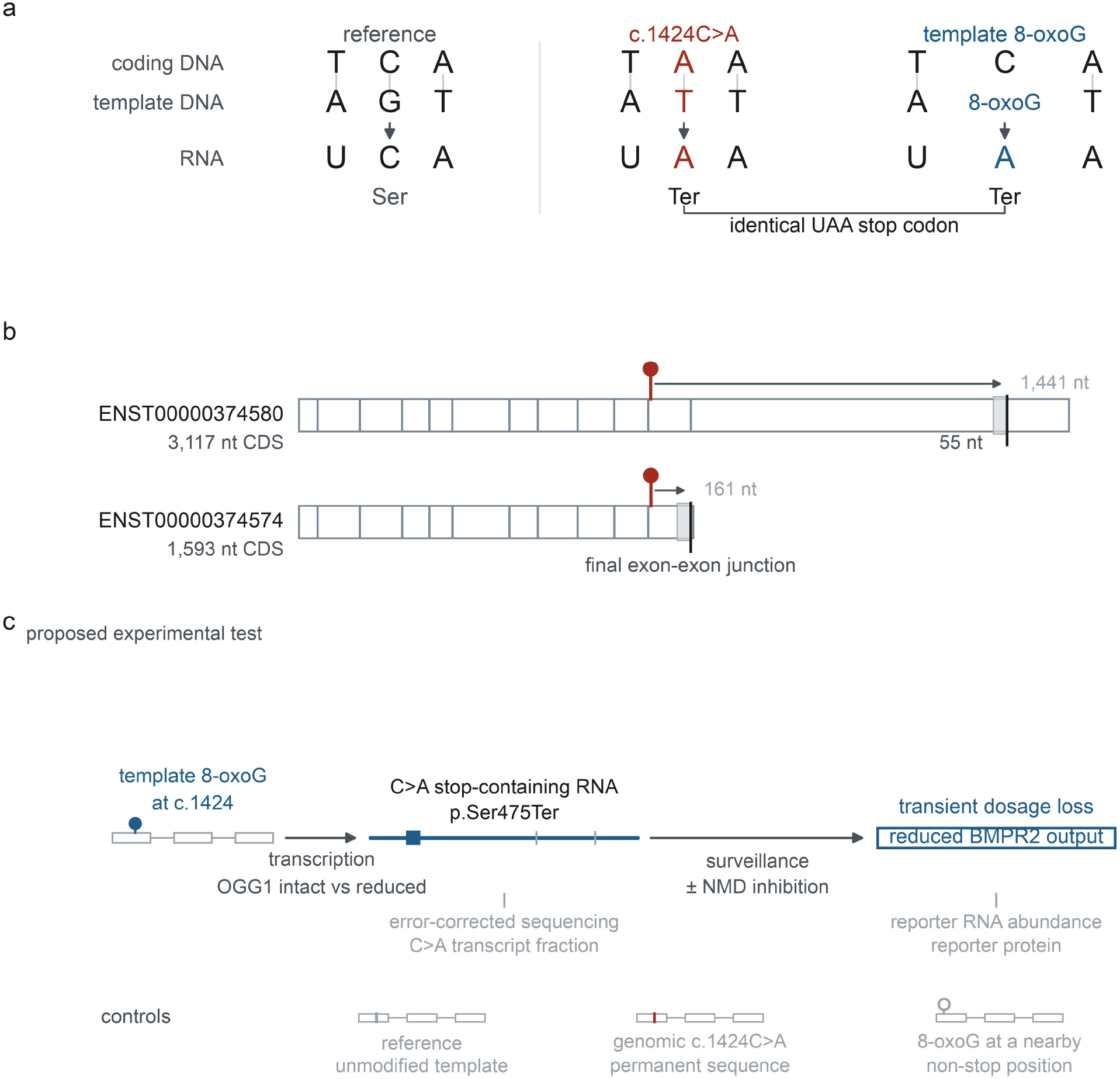
*BMPR2* c.1424 defines a direct experimental test of transient dosage loss. Red marks the permanent genomic route and blue the 8-oxoG-dependent route. (**a**) The pathogenic c.1424C>A allele and a template-strand 8-oxoG generate the same UAA-containing RNA at codon 475, although only the genomic route changes DNA. (**b**) p.Ser475Ter is predicted NMD-prone in both GENCODE-primary *BMPR2* transcripts containing the site. (**c**) Proposed experimental test, labeled as such on the panel, tracing a site-specific template lesion through C>A RNA, NMD surveillance, and *BMPR2* output. The permanent stop allele and a nearby non-stop lesion serve as sequence and lesion controls. NMD denotes nonsense-mediated decay, and Ter denotes translation termination.

## 3 Discussion

This analysis identifies defined positions where 8-oxoG transcriptional miscoding could generate the same premature-stop RNA as pathogenic PAH alleles without altering genomic DNA. Among 116 predicted NMD-prone positions in dosage-sensitive PAH genes, eight reproduce the RNA sequence generated by a ClinVar pathogenic or likely pathogenic stop variant. *BMPR2* c.1424 provides the strongest genetic precedent, because the corresponding genomic c.1424C>A allele is expert-panel reviewed and both routes are predicted to generate the same UAA-containing transcript. Human genetics establishes the pathogenicity of the genomic allele encoding this truncating transcript, while 8-oxoG miscoding would provide a transient route to the same RNA sequence.

The two routes differ in how long they act. A germline truncating allele continuously produces the altered transcript from one chromosome, whereas a lesion-driven error exists only while the damaged template remains unrepaired and affects only transcripts generated during that interval. The resulting reduction in gene output could be episodic and heterogeneous across pulmonary vascular cells, allowing a genetically intact cell to enter a temporary low-*BMPR2* state. Its magnitude would depend on lesion occupancy, transcription rate, miscoding probability, repair kinetics, RNA surveillance, and protein turnover. This mechanism is distinct from regulatory effects of genomic 8-oxoG that alter transcription without changing the RNA sequence [3,16], because the event considered here creates a different transcript.

The oxidative environment of the pulmonary hypertensive vasculature makes this mechanism biologically plausible. Severe pulmonary hypertension is associated with oxidative nucleic-acid damage [1], PAH endothelial cells show increased reactive-oxygen-species-associated DNA damage [2], and hypoxic signaling can generate sequence-localized oxidative guanine lesions in pulmonary artery endothelial DNA [3]. Circulating 8-hydroxy-2’-deoxyguanosine tracks disease severity in PAH despite not being globally elevated across unselected patients [17]. These observations do not identify *BMPR2* c.1424 as an endogenous oxidation hotspot, but they place the proposed lesion-driven mechanism within a disease environment where oxidative DNA damage and dosage-sensitive gene loss coexist.

Several limitations define what the present analysis can establish. The NMD assignments are predictions from transcript architecture, depend in part on the start-proximal surveillance rule, and vary across transcript isoforms at some positions. The endogenous frequency of template-strand 8-oxoG at the nominated sites in pulmonary vascular cells is unknown; the available lesion maps were generated in nonvascular cells, and the coordinate-level support at seven of the eight exact-precedent sites comes from an assay that does not identify the modified strand. The pulmonary vascular expression filter derives from reference lung rather than PAH tissue. A transient miscoding event would affect only a fraction of the transcripts generated while the lesion persists, so the pathogenicity of a constitutive genomic truncating allele does not establish that low-frequency transient production is sufficient to reduce functional gene dosage. These limitations define the experimental questions rather than the sequence accessibility addressed here.

*BMPR2* c.1424 provides a direct experimental system for resolving these questions. A site-specific template 8-oxoG can be compared with the permanent c.1424C>A allele and appropriate lesion controls to determine whether the lesion generates the predicted C>A transcript above the background transcription-error rate [18], whether repair limits its persistence, whether NMD inhibition stabilizes the premature-stop transcript, and whether *BMPR2* RNA or protein output falls (Figure 3c).

Demonstrating this sequence of events would establish a mechanism in which repairable oxidative DNA damage transiently reproduces a loss-of-function transcript encoded by a pathogenic PAH allele without genomic mutation. More broadly, it would extend the consequences of oxidative DNA damage beyond persistent mutation and transcriptional regulation to the temporary generation of disease-associated RNA sequences from genetically intact alleles.

## 4 Methods

### Gene panel

The analysis included all 24 genes carrying a ClinGen curation for pulmonary arterial hypertension (MONDO:0015924), retrieved 2026-09-19. *We included EIF2AK4, ENG*, and *ACVRL1* in a broader pulmonary vascular panel to capture overlap biology with pulmonary veno-occlusive disease, capillary hemangiomatosis, and hereditary hemorrhagic telangiectasia, yielding 27 genes. *ENG* and *ACVRL1* carry no PAH curation and are not treated as PAH-curated genes. We assigned gene-validity classification and inheritance mode from the ClinGen framework [19]; we assigned the dominant mechanism from ClinGen curation and the primary disease-gene literature, retained both as annotations, and applied them only during downstream filtering. The rationale and source for each gene are provided in Supplementary Table S1.

### Transcript retrieval

Transcripts were retrieved from Ensembl and verified as identical to MANE Select v1.5 [20] for all 27 genes, with intact start and stop codons, reading frame, and agreement with the annotated protein. The Ensembl transcript identifier and MANE Select RefSeq accession for every gene are given in Supplementary Table S1, and the full sequence audit is in the Supplementary Methods.

### Enumeration and NMD classification

C>A was applied at every coding-strand cytosine; the codon was retranslated and the consequence recorded, excluding the native stop codon. Exon boundaries were expressed in coding coordinates, and the 55-nucleotide rule was applied relative to the last exon-exon junction. Throughout, the distance from a premature termination codon to the final exon-exon junction is defined as the number of coding nucleotides from the last base of that stop codon to the last coding base preceding the junction, and this single definition is used for classification, for every reported value and in every figure [13]. Stops within 200 coding nucleotides of the start codon were classified separately as predicted inefficient-surveillance positions. This start-proximal rule is a heuristic rather than a measured threshold. Both it and the final-junction distance were varied in the sensitivity analysis shown in Supplementary Table S2, spanning 50 or 55 nucleotides for the junction rule and 150, 200 or 250 nucleotides for the start-proximal exception. Stops were assigned in the following order: (i) single-exon transcript, (ii) downstream of the final exon-exon junction, (iii) within 55 nucleotides upstream of the final junction, (iv) within the first 200 coding nucleotides, and (v) otherwise predicted NMD-prone. Junction-based rules take precedence over the start-proximal rule.

### Expression

Per-cell-type expression for all 27 genes was retrieved from the integrated Human Lung Cell Atlas [21] via the CELLxGENE Discover gene-expression API. A gene entered the vascular-expressed subset if it was detected in at least 10% of cells in at least one lung vascular or vascular-associated population. The full population list and the core-vascular flag are provided in Supplementary Table S4, while expression-threshold sensitivity is reported in Supplementary Table S3. The strict PAH set remained unchanged at 116 positions and eight exact precedents under both the extended and core vascular definitions at the 5% and 10% detection thresholds. Values are derived from reference lung tissue, not from PAH pulmonary arterioles.

### ClinVar variants and strict candidate definition

ClinVar variant_summary.txt.gz (GRCh38 assembly rows) [22] was downloaded on 2026-09-20 and frozen before filtering. We then filtered it to the panel, HGVS-parsed it, and reconciled it against the retrieved coding sequences, using the match criteria given in the Supplementary methods. Every surveillance classification reported in this work is a prediction from transcript architecture rather than a measurement of RNA decay. A strict PAH candidate position is a C>A-accessible coding position that would create a premature termination codon predicted NMD-prone under the primary rule applied to the MANE Select transcript, lies in a gene with ClinGen Definitive autosomal-dominant PAH gene-disease validity and an established haploinsufficiency mechanism, and passes the prespecified pulmonary vascular-expression filter, giving 116 positions in four genes. The primary surveillance call is made in the MANE Select transcript, and these positions are not uniformly NMD-prone across isoforms: 51 are predicted NMD-prone in every GENCODE-primary transcript carrying the position, 8 differ between such transcripts, and 57 occur in a single primary transcript. ClinVar was queried only after the candidate set had been defined and was used for the retrospective genomic-precedent analysis alone. For the eight strict precedents, we read the asserted disease condition from each ClinVar record rather than inferring it from the gene, and all eight involve pulmonary arterial hypertension or primary pulmonary hypertension. Variation identifiers, review status, and the asserted condition for each are given in Supplementary Table S5. We computed the premature-stop denominator reported in Results separately from the archived monthly release *variant_summary_2026-09*.*txt*.*gz*, a dated release preceding the analysis snapshot, scanned for pathogenic and likely pathogenic premature-stop single-nucleotide variants in the four strict genes. We use two frozen releases because the candidate reconciliation and the denominator were computed at different times, and the archived release predates the analysis snapshot; the denominator therefore reflects the same database state as the exact precedents rather than a later state containing subsequently deposited variants. The per-variant result is deposited as *PREMATURE_STOP_DENOMINATOR*.*tsv*.

### Published lesion maps

Two primary published datasets were used as orthogonal annotations and were never combined into a composite score: endogenous CLAPS-seq coverage in HeLa-S3 cells [14] (GSE181312, hg19, three wild-type replicates, per-base coverage that is not strand-resolved) and strand-resolved click-code-seq calls in unexposed HAP1 cells [15] (GSE272366, GRCh38, three replicates, per-site calls). Coverage at each candidate coordinate was read from the deposited tracks after conversion to the assay build. Supplementary Methods provide strand handling, coordinate verification, and per-dataset properties, and Supplementary Table S5 lists per-site values for the eight positions with an exact precedent.

### Candidate reference

A machine-readable reference of all 17,880 sequence-accessible coding positions accompanies the manuscript as *PAH_8oxoG_CtoA_candidate_reference_GRCh38*.*tsv*.*gz* and includes the genomic coordinate, transcript, codon consequence, surveillance prediction, isoform robustness, and exact ClinVar precedent where present. Presence in the file indicates that a C>A transcriptional event is sequence-compatible at that coding position, not that the paired template guanine is oxidized. The archive also carries a BED representation of the same positions, the strict PAH candidate set, the sensitivity analyses, figure source data, the source manifest, and the analysis code.

## Data availability

All datasets are public. Variant records are from ClinVar, gene-disease validity classifications are from the ClinGen gene-validity export, lung expression is from the CELLxGENE Discover integrated lung corpus, and transcript and protein annotations are from Ensembl and MANE. The published oxidized-guanine maps are GSE181312 (CLAPS-seq) and GSE272366 (click-code-seq). The candidate reference of all 17,880 sequence-accessible coding positions accompanies this manuscript. The source manifest in that archive records the accession, retrieval date, and genome build for every input, together with a checksum and byte size for the one input that is frozen as a file in the release. Code and derived data are available at https://github.com/Marciano-Lab/8oxoG-PAH-transcriptional-mutagenesis.

## Code availability

The analysis code that generates every submitted table and figure accompanies this manuscript and runs from the archived derived inputs without network access. The archive will be deposited in a public repository under a citable digital object identifier at publication, and the corresponding release tag and commit will be given here.

## Author contributions

All authors designed and performed the analyses. D.P.M. wrote the manuscript. All authors reviewed and revised the manuscript and approved the final version.

## Ethics approval and consent to participate

Not applicable. The study analyzed only publicly available, de-identified datasets and generated no new human or animal data.

## Consent for publication

Not applicable.

