## Supplemental_Information for "8-Oxoguanine transcriptional miscoding can transiently reproduce PAH-associated premature-stop transcripts"

### Supplementary methods

The procedures below support the analyses reported in the main text. They are recorded here in full so that the main Methods can state what was done without reproducing every verification step. The accompanying archive contains the exact analysis inputs, derived outputs and verification records used for the submitted results.

**Coding-sequence audit.** Every analyzed coding sequence passed six checks: an ATG start codon, a canonical stop codon, a length divisible by three, no internal stop codon, agreement between the summed exon lengths and the sequence length, and agreement with the annotated protein length. The C>A enumeration was additionally verified against a brute-force pass over all 64 codons.

**Variant reconciliation criteria.** Each ClinVar record retained for the panel was reconciled base by base and residue by residue against the retrieved coding sequence. A match required an identical coding position, reference base, alternate base and resulting amino-acid consequence, and records failing any of these were discarded rather than corrected.

**Expression filter in full.** Per-cell-type expression for all 27 genes was retrieved from the integrated Human Lung Cell Atlas [21] via the CELLxGENE Discover gene-expression API, covering 35 annotated lung vascular and vascular-associated populations including pulmonary artery endothelial cells, smooth muscle cells of the pulmonary artery, lung pericytes and lung fibroblasts. Expression is reported as the fraction of cells in which the gene is detected, aggregated with cell-type rollup. A gene entered the vascular-expressed subset if detected in at least 10% of cells in at least one of these populations. For the expression criterion, the main specification used the extended vascular-associated cell set at a 10% detection threshold, because pulmonary vascular remodeling involves endothelial, smooth-muscle, mural and perivascular stromal compartments. The main disease-focused analysis reported in the Results is the strict PAH set. Of the 35 populations, 23 form the narrower core vascular subset (14 vascular endothelial, 7 smooth-muscle and 2 mural or pericyte populations). The remaining 12 are perivascular stromal or lymphatic endothelial and are described as vascular-associated rather than vascular (Supplementary Table S4). The core subset and alternative detection thresholds were evaluated in sensitivity analyses, which left the strict PAH candidate set unchanged at 116 positions and eight exact precedents under both the extended and core vascular definitions at the 5% and 10% detection thresholds. Values derive from reference lung tissue, not from PAH pulmonary arterioles.

**Strand handling and coordinate verification.** The modified base sits on the template strand, so every modification-side quantity is computed there. Coding sequences were reverse-complemented to give the template strand in 5'-to-3' orientation, and per-base values were reindexed to coding coordinates. Coding-to-genomic coordinates were reconstructed from coding-clipped exon blocks and exhaustively verified against GRCh38: each block was re-fetched from the reference, reverse-complemented for minus-strand genes, reassembled in coding order and compared position by position with the MANE sequence. All 66,183 coding bases, including all 17,880 C>A-accessible cytosines, were concordant.

**Strict PAH disease-specific analysis.** The strict filter is defined in the main Methods. *ENG* and *ACVRL1* have no PAH curation and enter the wider pulmonary vascular panel through PAH-overlapping hereditary hemorrhagic telangiectasia. *SMAD4* is curated for PAH as Disputed. All downstream steps were recomputed unchanged on the strict panel.

**Transcript-isoform robustness.** All protein-coding transcripts of each contributing gene were retrieved from Ensembl and restricted to GENCODE-primary annotations, giving 30 transcripts. For each transcript a coding-position index was constructed from its coding exon blocks in transcript orientation, and each strict PAH candidate position was mapped through its genomic coordinate. The same architectural rules were applied per transcript. A position was classed NMD-robust when predicted NMD-prone in every GENCODE-primary transcript containing it, NMD-variable when predicted to escape in at least one, and one primary transcript only when contained in a single primary transcript.

**Data versions.** The accession, version, retrieval date and, where the input is frozen as a file, the checksum and byte size of every external input are recorded in *SOURCE_MANIFEST.tsv*. CELLxGENE Discover whole-corpus gene expression from the integrated Human Lung Cell Atlas was queried 2026-09-20 (snapshot 1762972271). Coding sequences, exon structures and transcript tags were retrieved from Ensembl release 116 on 2026-09-19, and the analyzed transcript for each of the 27 genes is the MANE Select transcript, verified by the Ensembl MANE Select tag and by exact coding-sequence identity to the paired RefSeq record.

**Candidate reference file.** For every gene the reconstructed coding sequence, read base by base from the reference genome through the resulting coordinate map and reverse-complemented for minus-strand genes, was verified identical to the analysis transcript, and every reference base in the released file was confirmed against the reference genome.

### Supplementary results

**Published lesion-map annotation.** Every accessible coding position was assigned a genomic coordinate from its coding-clipped exon blocks, and the modified guanine was placed on the strand opposite the coding strand. Coordinate mapping was first spot-checked against the GRCh38 reference on 60 sampled positions, all concordant, which tested the coding-to-genomic conversion including strand handling. The reference base at every one of the 17,880 accessible positions in the released coordinate file was then confirmed against the reference genome, so the exhaustive check supersedes the sample for that transformation. Two primary published lesion-map datasets were used as orthogonal annotations and were never combined into a composite score: endogenous CLAPS-seq coverage in HeLa-S3 (GSE181312, hg19, three wild-type replicates) and strand-resolved click-code-seq calls in unexposed HAP1 cells (GSE272366, GRCh38, three replicates). Candidates were lifted to hg19 for CLAPS-seq and every candidate position's hg19 reference base was confirmed. Exact-coordinate overlap was the primary analysis, and map eligibility was defined before testing.

**Published oxidative-lesion maps.** The two primary datasets differ in what they can establish. CLAPS-seq is per-base but not strand resolved, and its files are RPKM-normalized coverage tracks rather than called lesion sites, so nonzero coverage at a coordinate cannot establish that the modified guanine lies on the transcribed strand. Seven of the eight exact-precedent coordinates carry nonzero coverage, including *BMPR2* c.1424 in two of three wild-type replicates. Click-code-seq [15] is the only strand-resolved single-nucleotide dataset used, and no position with an exact precedent carries a template-strand oxidized-guanine call in it. As assay context, one other strict PAH candidate position, *TBX4* c.690, does carry such a call in one of three replicates. Both resources derive from nonvascular cells, and all strict PAH candidate positions were map-eligible.

### Supplementary figures

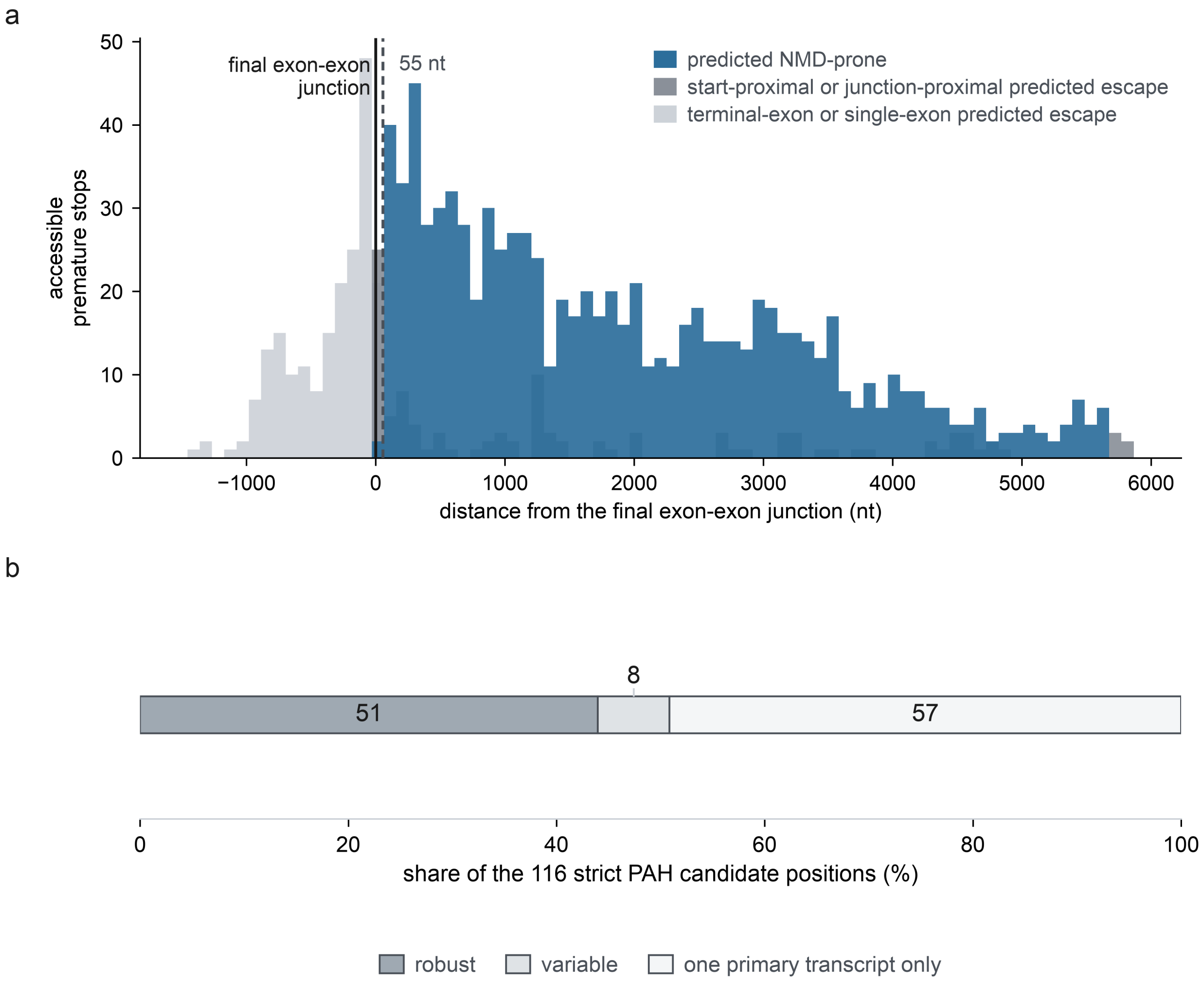


**Supplementary Figure S1. Transcript architecture of accessible premature stops.** (**a**) Distance of every accessible premature stop from the final exon-exon junction. The solid line marks the junction, at which the distance is zero, and the dashed line marks the 55-nucleotide junction-proximal window. Positive values lie upstream of the junction and negative values lie downstream of it. Bars are colored by architectural class: predicted NMD-prone, predicted escape in the terminal exon or a single-exon coding sequence, and the two boundary-based escape classes taken together, one of which is defined by distance from the start codon and so appears anywhere on this axis. (**b**) Predicted surveillance behavior of the strict PAH positions across the GENCODE-primary transcripts containing each position, classified as robust, variable or one primary transcript only. Class definitions are given in the Methods. NMD denotes nonsense-mediated decay.

### Supplementary tables

The five tables below are the publication-facing Supplementary Tables. Each is rendered here directly from the tab-separated file of the same name in the accompanying archive, so the printed table and the deposited file cannot diverge. Where a table carries more columns than fit the page, the printed view is a subset and the deposited file holds the full column set.

**Supplementary Table S1.** The 27-gene pulmonary vascular panel with ClinGen gene-disease validity for pulmonary arterial hypertension, inheritance mode, established disease mechanism and its source, the analyzed Ensembl MANE Select transcript and its RefSeq accession, reference-lung expression, accessible premature-stop and predicted NMD-prone counts, and the number of strict PAH candidate positions each gene contributes. Inheritance is abbreviated AD for autosomal dominant, AR for autosomal recessive, SD for semidominant and UD for undetermined, and mechanism is abbreviated HI for haploinsufficiency and LoF for loss of function. The haploinsufficiency mechanism for *BMPR2*, *KDR*, *SMAD9* and *TBX4*, the four genes contributing strict candidate positions, is taken from the ClinGen pulmonary arterial hypertension gene-disease validity curation. Per-gene mechanism sources for the remaining genes, several of which cite primary literature, and the per-cell-type expression values are in the deposited file *Table_S1.tsv*.

| Gene | ClinGen PAH validity | Inheritance | Mechanism | MANE RefSeq | Max vascular % | Accessible stops | NMD-prone | Strict |
|---|---|---|---|---|---|---|---|---|
| ABCC8 | Moderate | AD | mixed LoF/mutant | NM_000352.6 | 8.3 | 67 | 63 | 0 |
| ACVRL1 | not curated for PAH | AD | HI | NM_000020.3 | 50.2 | 28 | 21 | 0 |
| AQP1 | Limited | AD | unclear | NM_198098.4 | 77.1 | 12 | 9 | 0 |
| ATP13A3 | Definitive | SD | mixed LoF/mutant | NM_001367549.1 | 38.7 | 48 | 41 | 0 |
| BMP10 | Limited | AD | HI | NM_014482.3 | 0.0 | 21 | 3 | 0 |
| BMPR1A | Disputed | UD | HI | NM_004329.3 | 33.1 | 34 | 25 | 0 |
| BMPR1B | Disputed | UD | unclear | NM_001203.3 | 13.7 | 25 | 17 | 0 |
| BMPR2 | Definitive | AD | HI | NM_001204.7 | 81.6 | 38 | 27 | 27 |
| CAV1 | Definitive | AD | mixed LoF/mutant | NM_001753.5 | 92.7 | 11 | 0 | 0 |
| EIF2AK4 | not curated for PAH | AR | recessive | NM_001013703.4 | 10.7 | 70 | 62 | 0 |
| ENG | not curated for PAH | AD | HI | NM_001114753.3 | 82.7 | 32 | 24 | 0 |
| FBLN2 | Limited | AD | unclear | NM_001004019.2 | 44.8 | 108 | 93 | 0 |
| GDF2 | Definitive | AD | HI | NM_016204.4 | 0.1 | 14 | 4 | 0 |
| GGCX | Moderate | AD | recessive | NM_000821.7 | 5.1 | 36 | 28 | 0 |
| KCNK3 | Definitive | AD | mixed LoF/mutant | NM_002246.3 | 42.0 | 36 | 0 | 0 |
| KDR | Definitive | AD | HI | NM_002253.4 | 31.4 | 62 | 55 | 55 |
| KLF2 | Limited | AD | unclear | NM_016270.4 | 62.6 | 21 | 9 | 0 |
| KLK1 | Limited | AD | unclear | NM_002257.4 | 0.7 | 10 | 4 | 0 |
| NOTCH3 | Disputed | AD | mutant protein | NM_000435.3 | 55.6 | 222 | 206 | 0 |
| PDGFD | Limited | AD | unclear | NM_025208.5 | 25.0 | 26 | 16 | 0 |
| SMAD1 | Disputed | AD | unclear | NM_005900.3 | 37.6 | 22 | 17 | 0 |
| SMAD4 | Disputed | AD | HI | NM_005359.6 | 15.4 | 26 | 20 | 0 |
| SMAD9 | Definitive | AD | HI | NM_001127217.3 | 14.2 | 28 | 25 | 25 |
| SOX17 | Definitive | AD | HI | NM_022454.4 | 40.5 | 25 | 0 | 0 |
| TBX4 | Definitive | AD | HI | NM_001321120.2 | 22.8 | 29 | 9 | 9 |
| TET2 | Moderate | AD | HI | NM_001127208.3 | 14.6 | 86 | 60 | 0 |
| TOPBP1 | No Known Disease Relationship | AD | none | NM_007027.4 | 7.5 | 55 | 45 | 0 |

**Supplementary Table S2.** Sensitivity of the strict PAH candidate set to the surveillance rule. The junction and start-proximal thresholds were varied around the primary specification of 55 and 200 nucleotides. Deposited as *Table_S2.tsv*.

| Junction rule (nt) | Start rule (nt) | Panel NMD-prone | Strict positions | Exact precedents | BMPR2 precedents |
|---|---|---|---|---|---|
| 50 | 150 | 909 | 121 | 8 | 5 |
| 50 | 200 | 885 | 116 | 8 | 5 |
| 50 | 250 | 862 | 113 | 7 | 5 |
| 55 | 150 | 907 | 121 | 8 | 5 |
| 55 | 200 | 883 | 116 | 8 | 5 |
| 55 | 250 | 860 | 113 | 7 | 5 |

**Supplementary Table S3.** Sensitivity of the strict PAH candidate set to the reference-lung expression filter across detection thresholds and both cell-type definitions. Deposited as *Table_S3.tsv*.

| Cell-type set | Threshold % | Strict positions | Exact precedents | Genes contributing |
|---|---|---|---|---|
| extended vascular-associated (all 35 populations) | 5 | 116 | 8 | 4 |

| Cell-type set | Threshold % | Strict positions | Exact precedents | Genes contributing |
|---|---|---|---|---|
| extended vascular-associated (all 35 populations) | 10 | 116 | 8 | 4 |
| extended vascular-associated (all 35 populations) | 20 | 91 | 5 | 3 |
| core vascular (endothelium, smooth muscle, pericytes) | 5 | 116 | 8 | 4 |
| core vascular (endothelium, smooth muscle, pericytes) | 10 | 116 | 8 | 4 |
| core vascular (endothelium, smooth muscle, pericytes) | 20 | 82 | 5 | 2 |

**Supplementary Table S4.** The 35 lung vascular and vascular-associated populations used for the expression filter, including Cell Ontology identifiers, compartment assignment and the core-vascular flag. Machine-readable only, provided as *Table_S4.tsv*.

**Supplementary Table S5.** The eight strict PAH candidate positions whose predicted C>A RNA sequence exactly matches a pathogenic or likely pathogenic genomic premature-stop allele, with the ClinVar record, the asserted disease condition read from that record, the number of conditions the record carries, the junction distance in the MANE Select transcript, isoform robustness and the per-site lesion-map values. Genomic and template coordinates, CLAPS-seq replicate support and mean signal, the click-code-seq template call, both assay builds and the assay limitation are in the deposited file *Table_S5.tsv*. The *SMAD9* c.204C>A record carries two asserted conditions, the pulmonary arterial hypertension entry shown here and a second submitter entry of *not provided*, which is carried in the deposited file.

| Gene | cDNA | Protein | Variation ID | Classification | Review status | Asserted condition | Junction distance (nt) | Isoform robustness |
|---|---|---|---|---|---|---|---|---|
| BMPR2 | c.320C>A | p.Ser107Ter | 1453966 | Pathogenic | criteria provided, single submitter | Primary pulmonary hypertension | 2545 | NMD-variable |
| BMPR2 | c.339C>A | p.Tyr113Ter | 425761 | Pathogenic | no assertion criteria provided | Pulmonary hypertension, primary, 1 | 2527 | NMD-variable |
| BMPR2 | c.507C>A | p.Cys169Ter | 8804 | Pathogenic | no assertion criteria provided | Pulmonary hypertension, primary, 1 | 2359 | NMD-robust |
| BMPR2 | c.864C>A | p.Cys288Ter | 4716557 | Pathogenic | criteria provided, single submitter | Primary pulmonary hypertension | 2002 | NMD-robust |
| BMPR2 | c.1424C>A | p.Ser475Ter | 425943 | Pathogenic | reviewed by expert panel | Pulmonary arterial hypertension | 1441 | NMD-robust |
| SMAD9 | c.204C>A | p.Cys68Ter | 548690 | Pathogenic/Likely pathogenic | criteria provided, multiple submitters, no conflicts | Pulmonary arterial hypertension associated with congenital heart disease | 1056 | NMD-robust |
| SMAD9 | c.606C>A | p.Cys202Ter | 6515 | Pathogenic | no assertion criteria provided | Pulmonary hypertension, primary, 2 | 654 | NMD-robust |
| SMAD9 | c.801C>A | p.Tyr267Ter | 4077556 | Likely pathogenic | criteria provided, single submitter | Pulmonary hypertension, primary, 2 | 459 | NMD-robust |

**Supplementary files**

The bioRxiv Supplementary Data archive contains Supplementary Tables S1 to S5, the complete 17,880-position candidate reference and its BED representation, the 116-position strict candidate set and source data for each data-bearing figure panel, listed in *BIORXIV_SUPPLEMENTARY_DATA_INDEX.tsv* with a one-line description, row and column counts and a SHA-256 checksum for each file, and with every column defined in *DATA_DICTIONARY.tsv*. The complete analysis code, frozen derived inputs, provenance records and computational environment are provided separately in the public repository release described under Code availability.

**Supplementary Tables S1 to S5** as tab-separated files, identical to the tables rendered above.

***PAH_8oxoG_CtoA_candidate_reference_GRCh38.tsv.gz*** the complete reference of all 17,880 C>A-accessible coding positions across the 27 panel genes, with codon consequence, predicted surveillance class, junction distance and the strict-set flag.

***PAH_8oxoG_CtoA_candidate_reference_GRCh38.bed.gz*** the same positions as BED6+4 on GRCh38, with 0-based half-open coordinates for genome-browser and interval-tool use.

***strict_pah_candidate_positions.tsv*** the 116 strict PAH candidate positions as a standalone table.

**Figure source data** one file per data-bearing figure panel, with a panel manifest that records which panels are schematics carrying no numeric source.

**Analysis code and provenance** the enumeration and figure scripts, the frozen input record with retrieval dates and checksums, and the environment specification.
